# Influence of miR-520e-mediated MAPK signaling pathway on HBV replication and regulation of hepatocellular carcinoma cells via targeting EphA2

**DOI:** 10.1101/341511

**Authors:** Jing-hui Tian, Wen-dong Liu, Zhi-yong Zhang, Li-hua Tang, Dong Li, Zhao-ju Tian, Shao-wei Lin, Ying-jie Li

**Affiliations:** Central Laboratory, Shandong Provincial Hospital Affiliated to Shandong University, Ji’nan 250021, China.; School of Public Health, Taishan Medical University, Taian 271016, China.; Department of Blood Transfusion, Affiliated Hospital of Weifang Medical University, Weifang 261031, China.; Clinical Laboratory, Dezhou People’s Hospital, Dezhou 253014, China.; Department of Blood Transfusion, Tai’an City Central Hospital, Tai’an 271000, China.; Department of Health Examination, Qilu Hospital, Shandong University, Jinan 250012, China.

**Keywords:** miR-520e, EphA2, Hepatocellular carcinoma

## Abstract

This paper aims to determine the role of miR-520e in the replication of hepatitis B virus (HBV) and the growth of hepatocellular carcinoma (HCC) cells. MiR-520e and EphA2 in HBV-positive HCC tissues and cells were detected. HepG2.2.15 and Huh7 cells transfected with pHBV1.2 were divided into Mock, NC, miR-520e mimic, miR-520e inhibitor, si-EphA2, and miR-520e inhibitor + si-EphA2 groups. MiR-520e, HBV DNA content, HBsAg and HBeAg levels, cell proliferation, apoptosis and protein expression of EphA2 and MAPK pathways were evaluated. Furthermore, rAAV81.3HBV infected-mouse model was established to detect HBV-DNA levels. MiR-520e was up-regulated and EphA2 was down-regulated in HBV-positive HCC tissues and cells (HepG2.2.15 and HepAD38). MiR-520e was decreased in Huh7-X and HepG2-X cells in which HBx was stably expressed, but miR-520e was dose-dependently elevated in Huh7-X, HepG2-X, and HepG2.2.15 cells after interfering HBx. Additionally, miR-520e mimic and si-EphA2 groups were apparently reduced in HBV DNA content, HBsAg and HBeAg levels, cell proliferation, and were enhanced in the expressions of EphA2, MAPK pathways and cell apoptosis. Furthermore, si-EphA2 can reverse the promotion effect of miR-520e inhibitor on the HBV replication and tumor cell growth Up-regulating miR-520e in rAAV81.3HBV infected-mouse resulted in the reduced EphA2 in liver tissues and HBV DNA content in serum. MiR-520e was found to be decreased in HBV-positive HCC tissues and cells, while over-expression of miR-520e blocked MAPK pathways via inhibiting EphA2, ultimately reducing HBV replication and inhibiting tumor cell growth.

## Introduction

Hepatocellular carcinoma (HCC) is believed as one of the most common tumors in the digestive system, which ranks the fifth most frequent cancer and the third leading cause of cancer-related death respectively all over the world (Lodato et al., 2006), but is usually diagnosed at the late or advanced stage due to the lack of early diagnosis solution, resulted in a poor prognosis (Boyault et al., 2007). As reported previously, hepatitis B virus (HBV) infection is considered as a major threat for HCC, and most patients suffered from HCC as a consequence of persistent HBV infection, especially in China (Hu et al., 2012). Moreover, over 350 million people was estimated to be chronically infected with HBV (Zhao et al., 2014), included 50% of male and 14% of female carriers, who would eventually die of liver cirrhosis and HCC (Ooi and Teoh, 2012). Thus, it is of great values to elucidate the pathogenesis of HBV infection-induced HCC.

MicroRNAs (miRNAs) are small non-coding RNA molecules, consisting of 19-25 nucleotides (nts), which can regulate gene expressions at the post transcriptional level by directly degrading mRNA or inhibiting its translation (Hu et al., 2017). Several lines of evidence demonstrated that miRNAs, as one of the crucial regulators, could mediate the interaction between the virus and the host (Li et al., 2016a; Umbach and Cullen, 2009). Besides, hepatitis B virus X protein (HBx) is well-known as the essential component for HBV replication, could induce the abnormal expression of miRNA. For example, HBx was suggested to increase miRNA-29a and miRNA-21 (Hou and Quan, 2017; Kong et al., 2011), but decrease miR-148, miR-152 and let-7 (Huang et al., 2010; Wang et al., 2010; Xu et al., 2013), to be involved in the pathogenesis of HBV-related HCC. In turn, miRNA could also regulate the replication of HBV. For instance, both miR-199a-3p and miR-210 have been found to play important roles in controlling HBV replication in HBV-positive HepG22.2.15 cells (Zhang et al., 2010). Besides, miR-15a/miR-16-1 was also identified to influence the growth of HCC cells via blocking the replication of HBV (Wang et al., 2013). To date, there was evidence indicated that miR-520 family, served as anti-onco-miRNAs, was implicated in regulating the tumorigenesis and development of various solid cancers (Yan et al., 2015). To be specific, miR-520b, as a member of miR-520 family, was reduced, while its target gene (*HBXIP*) was elevated, both of which were mediated by HBx, contributing to hepatocarcinogenesis *in vitro* and *in vivo* (Zhang et al., 2014). Of note, as another member of miR-520 family, miR-520e was also reported able to target NIK to mediate NIK/p-ERK1/2/NF-κB pathway, and thereby inducing the occurrence of HCC (Zhang et al., 2012). Hence, we hypothesized that miR-520e may exert functions in HBV-related HCC.

Furthermore, using the target gene prediction website, we found that EphA2 was the target gene of miR-520e, and it could regulate the MAPK signaling pathway in HCC cells, as indicated by Jin *et al.* (Jin et al., 2015). More importantly, HBx has been demonstrated to participate in the development and progression of HBV-related HCC by activating the MAPK signaling pathway (Tarn et al., 2001). Therefore, this study was undertaken to investigate whether miR-520e could regulate HBV replication and HCC growth by mediating the MAPK signaling pathway through targeting EphA2, hoping to provide new theoretical support for the diagnosis and treatment of HBV-related HCC.

## Materials and methods

### Ethics statement

This study was approved by the Ethics Committee of our hospital after discussion, which was in accordance with the ethics standards laid down in the 1964 Declaration of Helsinki, and all patients were informed of the purpose and content of the study and agreed to sign the informed consent.

### Tissue specimens

From October 2016 to October 2017, tumor tissues and adjacent tissues (> 2 cm) were collected from 45 HBV-positive HCC patients at the early stage who received hepatic carcinectomy in the Department of Hepatobiliary Surgery in our hospital. There were 32 male and 13 female patients with the age of 53.56 ± 10.45 years old. All tissue samples were verified by clinical and pathological examinations and then cryopreserved at -80°C.

### Cell lines and cell culture

HBV-positive cell lines (HepG2.2.15 and HepAd38), and HBV-negative cell lines (HepG2 and Huh7), were purchased from the American Type Culture Collection (ATCC). HepG2.2.15 (HepG2 cells integrated with 4 copies of HBV gene) can stably express HBV gene and secrete HBV virus particles to complete the replication of HBV, were cultured in MEM complete medium with 10% fetal bovine serum (FBS) and 380 μg/ml G418 and digested by 0.25% trypsin once every three days for sub-culturing. HepG2, HepAd38 and Huh7 cells were cultured in RPMI1640 complete medium containing 10% FBS in an incubator (at 37°C with 5% CO_2_) and digested by 0.25% trypsin once every two days for sub-culturing.

### Transfection of HCC cells with stably expressed HBx

Huh7-X cells, HCC cell line with stable expression of HBx, and HepG2-X cells were established and preserved in advance in the laboratory. To investigate the mechanism of miR-520e down-regulation, Huh7-X, HepG2-X and HepG2.2.15 cells were seeded to 6-well plates by 1×10^6^ cells/well and cell confluence reached 70-80% on the next day. The transfection was carried out according to the instructions on the LipofectamineTM2000 kit (Invitrogen). Cells were transfected with si-HBx plasmids by 0, 0.5, 1, 1.5 μg in each well, and the expression levels of HBx and miR-520e were detected after 48 h transfection.

### Dual-luciferase report gene assay

The miRNA target gene prediction software (microRNA.org) was used to predict the binding site of miR-520e and EphA2 3’UTR. We established the wild-type plasmid EphA2 WT and the mutant-type plasmid EphA2 MUT The transfection was performed by following the instructions using Lipofectamine^TM^2000 reagent (Invitrogen). HepG2 cells (purchased from ATCC) were transfected and classified into four groups: miR-520e mimic+ EphA2-WT group, mimic NC+ EphA2-WT group, miR-520e mimics+ EphA2-MUT group, and mimic NC+ EphA2-MUT group. After 48 h transfection, the luciferase activity of each sample (Promega) was detected by dual-luciferase reporter gene assay.

### Cell grouping and transfection

In this experiment, HepG2.2.15 cells and Huh7 cells transfected with pHBV1.2 (a 1.2-fold full-length HBV genome expression plasmid) were selected and divided into six groups: Mock group (cells without transfection), NC group (cells transfected with miR-520e negative control plasmid), miR-520e mimic group (cells transfected with miR-520e mimic), miR-520e inhibitor group (cells transfected with miR-520e inhibitor), si-EphA2 group (cells transfected with si-EphA2), and miR-520e inhibitor+ si-EphA2 inhibitor group (cells co-transfected with si-EphA2 and miR-520e inhibitor). All sequences were synthesized by the Shanghai GenePharma Co., Ltd and cell transfection was performed by following the instructions on the Lipofectamine™2000 Kit (Invitrogen).

### Detection of HBV DNA content

The supernatant of each cell sample was collected and detected for HBV DNA with a HBV DNA kit (QIAGEN). Next, HBV DNA copies were quantified by qRT-PCR.according to the instructions of the kit. The reaction conditions included: 40 cycles of 37°C for 5min, 94°C for 2min, 95°C for 5s, and 60°C for 40s. Then, fluorescence signals were collected at 60°C. The results were obtained and analyzed to draw a standard curve for each experiment. The mean value of three experiments was regarded as the final result.

### Detection of HBsAg and HBeAg by ELISA

The cell culture medium was collected 48 h post-transfection, and centrifuged for 5 min at the rate of 5000 rpm to obtain the supernatant. Then, the HBsAg and HBeAg viral protein levels were determined by ELISA kits (Shanghai Kehua Bio-Engineering Co., Ltd.) in the supernatant in line with the instructions, and a microplate reader was used to detect the absorbance value (OD) of each well under the wavelength of 450 nm.

### Cell proliferation by CCK8 assay

The cell proliferation was detected by CCK8 kits (Dojindo Molecular Technologies, Inc., Japan). First, cells were inoculated onto the 96-well plate at 10^3^ cells/well (200 μl/well) with 5 duplicates as a set. At the same time, blank control group (only added with medium) was set for 24 h, 48 h and 72 h of incubation. Next, CCK8 kit was added into each well (by 1: 9 in CCK8 to medium) for 1-4 h of incubation at 37°C. The black control group was used for zero setting and the absorbance value (OD) of each well under the wavelength of 450 nm was obtained from the microplate reader. The experiment was repeated for three times respectively.

### Cell apoptosis by flow cytometry

Annexin V-FITC/PI reagent (Shanghai BogooBiotechnology, Co., Ltd) was used to detect cell apoptosis. Cells were collected in the logarithmic growth phase, digested by 0.25% trypsin, washed twice with PBS buffer, centrifuged and added with 195 μL Binding Buffer to suspend cells. Next, 5 μL Annexin V-FITC was added into the cell suspension, which was placed for a while and centrifuged for cell collection. Then, 190 μL binding buffer was added to suspend cells, followed by the addition of 10 μL propidium iodide (PI) for mixing at room temperature and without exposure to light. Cell apoptosis was determined by a flow cytometer and the experiment was conducted for three times independently.

### qRT-PCR

Total RNA was extracted from tissues and cells with the Trizol reagent (Invitrogen), determined for concentration after detection of OD260/280 ratio of each RNA sample with a UV spectrophotometer, and cryopreserved at -80°C for later use. The primers in this experiment were designed by using Primer 5.0 and synthesized by Shanghai GenePharma Pharmaceutical Technology Co., Ltd (Table 1). The specific ring-shaped primer for the target miRNA, designed by the ABI Company, was applied for the reverse transcription of RNA into cDNA. The PCR amplification reaction was performed on the ABI Step One Plus real-time PCR system with the Taqman micro RNA Kit (ABI Company). The reaction conditions were as follows: pre-denaturation at 95°C for 10 min and 40 cycles of 95°C for 15 s and 60°C for 1 min. With U6 as the internal reference gene for miR-520e and β-actin for EphA2, the relative expression levels of target genes were calculated according to the 2^-ΔΔ^Ct method.

**Table 1.**
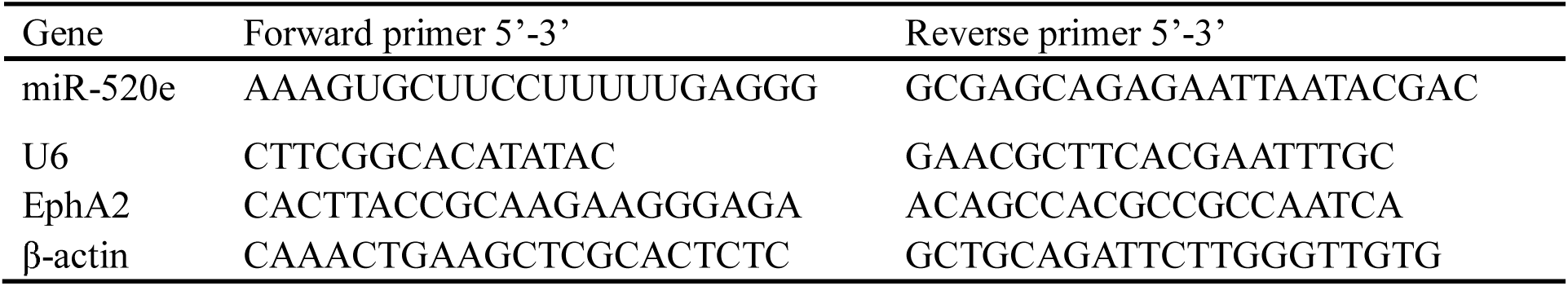
The Primer sequences for qRT-PCR applied in the current study

### Western blot

The proteins were extracted and added with loading buffer for 10 min of heating at 95°C with 30 μg/well. Next, 10% polyacrylamide gel electrophoresis was used to separate proteins, followed by wet transferring to the PVDF (polyvinylidene difluoride) membrane. Then, the membrane was placed in 5% bovine serum albumin (BSA) for blocking before the addition of primary antibodies for overnight reaction at 4°C at 1:1000 dilution, including EphA2, HBx, p38MAPK, p-p38MAPK, ERK1/2 and p-ERK1/2 (Cell Signaling Technology). After that, the membrane was washed with TBST (tris-buffered saline with Tween 20) for 3 times/5 min and corresponding secondary antibodies were added for 1 h of incubation at room temperature. At last, the membrane was washed for another 3 times/5 min before the chemiluminescent detection. With β-actin as the internal reference, the gray value of target bands was analyzed with the software Image J.

### Mice model establishment with adenovirus mediated HBV replication

Based on a previous study (Dai et al., 2014), 6-week old male C57BL/6 mice were injected with rAAV8-1.3HBV (1×10^11^ vg/200 μl/mouse) via the tail vein to construct model mice with the chronic HBV infection, which were randomly divided into two groups with 8 mice in each group: HBV+ miR-520e mimic group and HBV+NC group, which were injected with miR-520e mimic and negative control plasmid respectively by the high-pressure hydrodynamic method. Another 8 mice without HBV injection were set as the Normal group. Three days after injection, those mice were killed to take their liver tissues and tail vein blood. As described above, the relative expression of miR-520e in liver tissues and HBV DNA copy number in serum were analyzed by qRT-PCR.

### Statistical method

All experiments were repeated in three times. All data were analyzed by using the statistical software SPSS 21.0 (SPSS, Inc, Chicago, IL, USA), which were presented by mean ± standard deviation (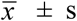). For difference between two groups, the data were compared by using the Student’s *t*-test, while among multiple groups were tested by One-Way ANOVA. *P* < 0.05 believed as the statistical significance.

## Results

### Expression of miR-520e and EphA2 in HBV-positive tissues and cell lines

The expression of miR-520e and EphA2 in HBV-positive HCC tissues and cell lines was detected by qRT-PCR. The results found that HBV-positive HCC tissues had significantly decreased miR-520e and apparently increased EphA2 compared to normal tissues (all *P* < 0.05). In addition, HBV-positive HepG2.2.15 and HepAd38 cells had lower levels of miR-520e than HBV-negative Huh7 and HepG2 cells, and HepAd38 cells showed the most significant reduction in miR-520e (all *P* < 0.05). Western blot was applied for the detection of EphA2 expression in cell lines, which demonstrated that EphA2 expression was higher in HepG2.2.15 and HepAd38 cells than Huh7 and HepG2 cells, and was the highest in HepAd38 cells (all *P* < 0.05, Figure 1).

**Figure 1.**
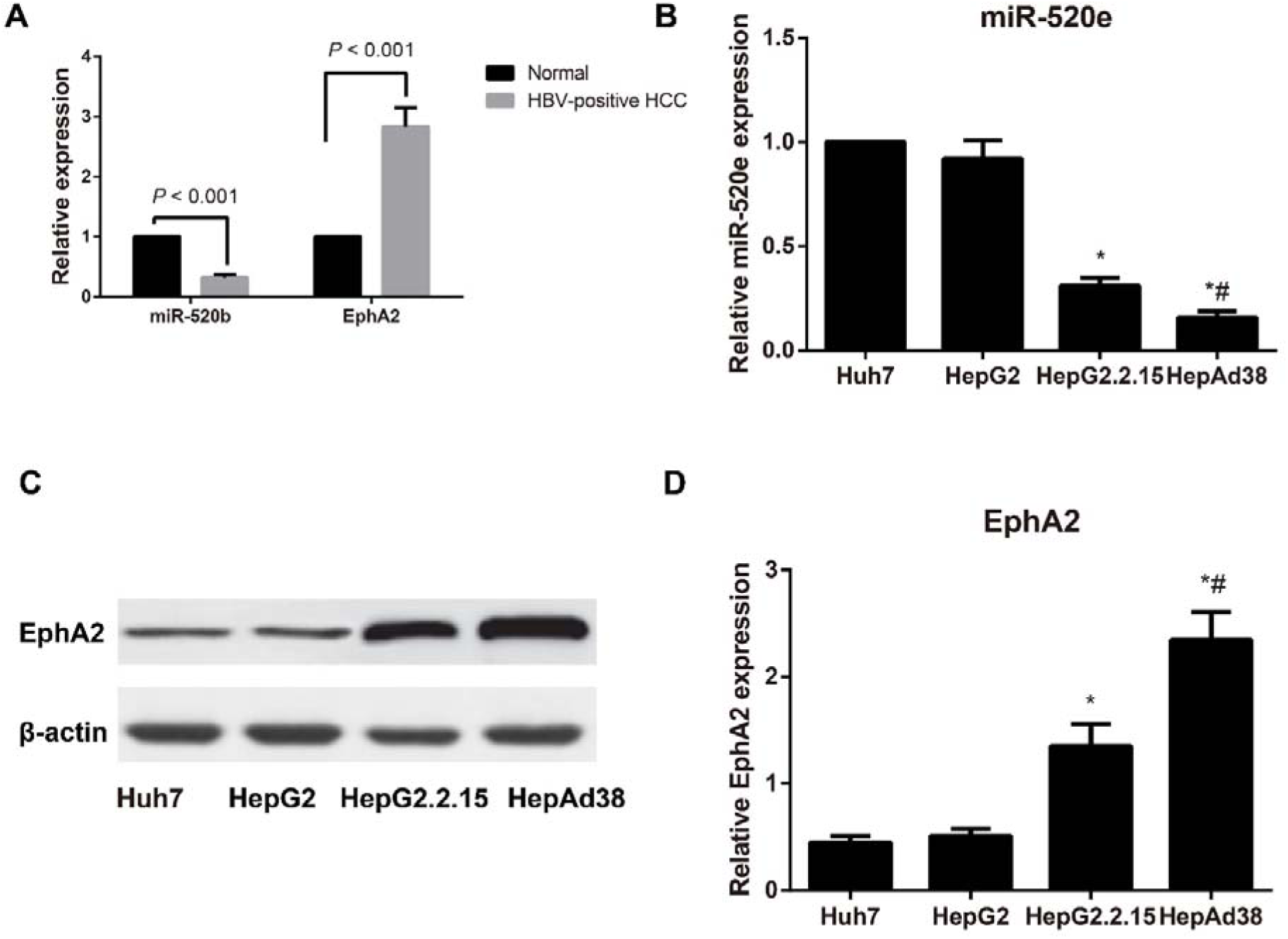
Expression of miR-520e and EphA2 in HBV-positive tissues and cell lines detected by qRT-PCR and Western blot. Note: A, Relative expression of miR-520e and EphA2 in HBV-positive tissues detected by qRT-PCR; B, Relative expression of miR-520e in HCC cell lines detected by qRT-PCR; C-D, Relative EphA2 expression in HCC cell lines detected by Western blot; *, *P* < 0.05 compared with Huh7 cells; ^#^, *P* < 0.05 compared with HepG2.2.15 cells.

### HBx down-regulates the expression of miR-520e

We constructed the Huh7-X and HepG2-X cell lines in which HBx was stably expressed were established, and we found the significant elevation of HBx protein in Huh7-X and HepG2-X cells (Figure 2A, B, all *P* < 0.05). In addition, the miR-520e levels presented to be remarkably lower in Huh7-X and HepG2-X cells than Huh7 and HepG2 cells (Figure 2C, *P* < 0.05). To further confirm the down-regulation of miR-520e by HBx, we found that the levels of miR-520e were dose-dependently enhanced via interfering HBx in a dose-dependent manner in Huh7-X, HepG2-X and HepG2.2.15 cells (Figure 2D, E), which suggested that HBx could reduce the expression of miR-520e (Figure 2F, *P* < 0.05).

**Figure 2.**
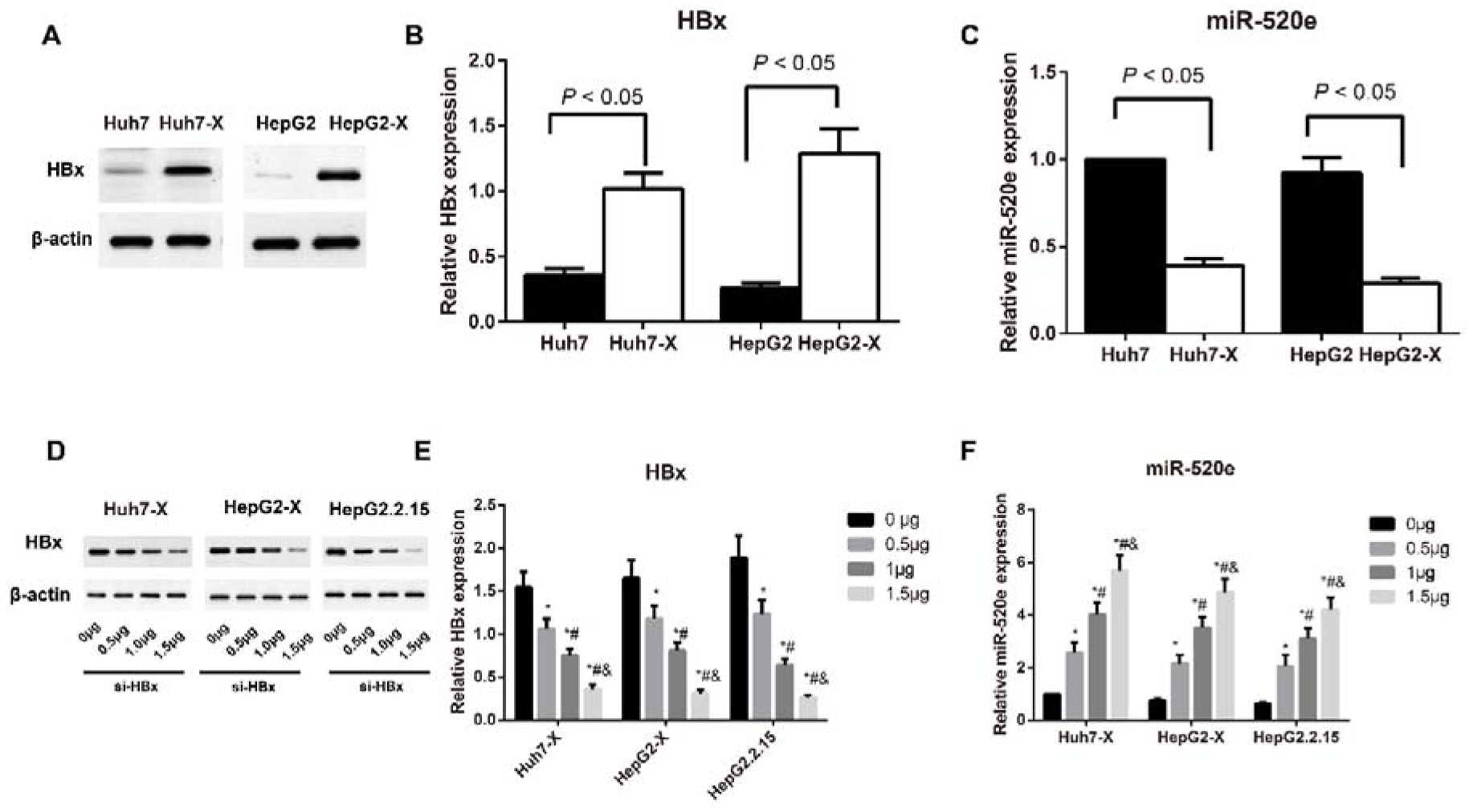
HBx can down-regulate the expression of miR-520e in HCC cells Note: A-B, Relative protein expression of HBx in Huh7-X and HepG2-X cells (in which HBx was stably expressed) detected by Western blot; C, The miR-520e levels in Huh7-X and HepG2-X cells (in which HBx was stably expressed) detected by qRT-PCR; D-E, Relative protein expression of HBx in Huh7-X, HepG2-X and HepG2.2.15 cells after interfering HBx detected by Western blot; F, The expression of miR-520e in Huh7-X, HepG2-X and HepG2.2.15 cells after interfering HBx detected by qRT-PCR; *, *P* < 0.05 compared with cells treated with 0 μg si-HBx; #, *P* < 0.05 compared with cells treated with 0.5 μg si-HBx; &, *P* < 0.05 compared with cells treated with 1 μg si-HBx.

### EphA2 is the target gene of miR-520e

Online miRNA target gene prediction website <u>(http://www.microrna.org)</u> was employed to predict the target gene of miR-520e, and we found EphA2 to be one of the target genes of miR-520e (Figure 3A). As illustrated by dual-luciferase reporter gene assay, the relative activity of luciferase was remarkably reduced after co-transfection with EphA2 3’UTR-WT and miR-520e mimic when compared with that of EphA2 3’UTR-WT + miR-520e NC group (*P* < 0.05). On the other hand, no change in luciferase activity was observed in EphA2 3’UTR -MUT+ miR-520e mimic group and EphA2 3’UTR -MUT + miR-520e NC group (all *P* > 0.05, Figure 3B), which suggested that EphA2 is the target gene of miR-520e.

**Figure 3.**
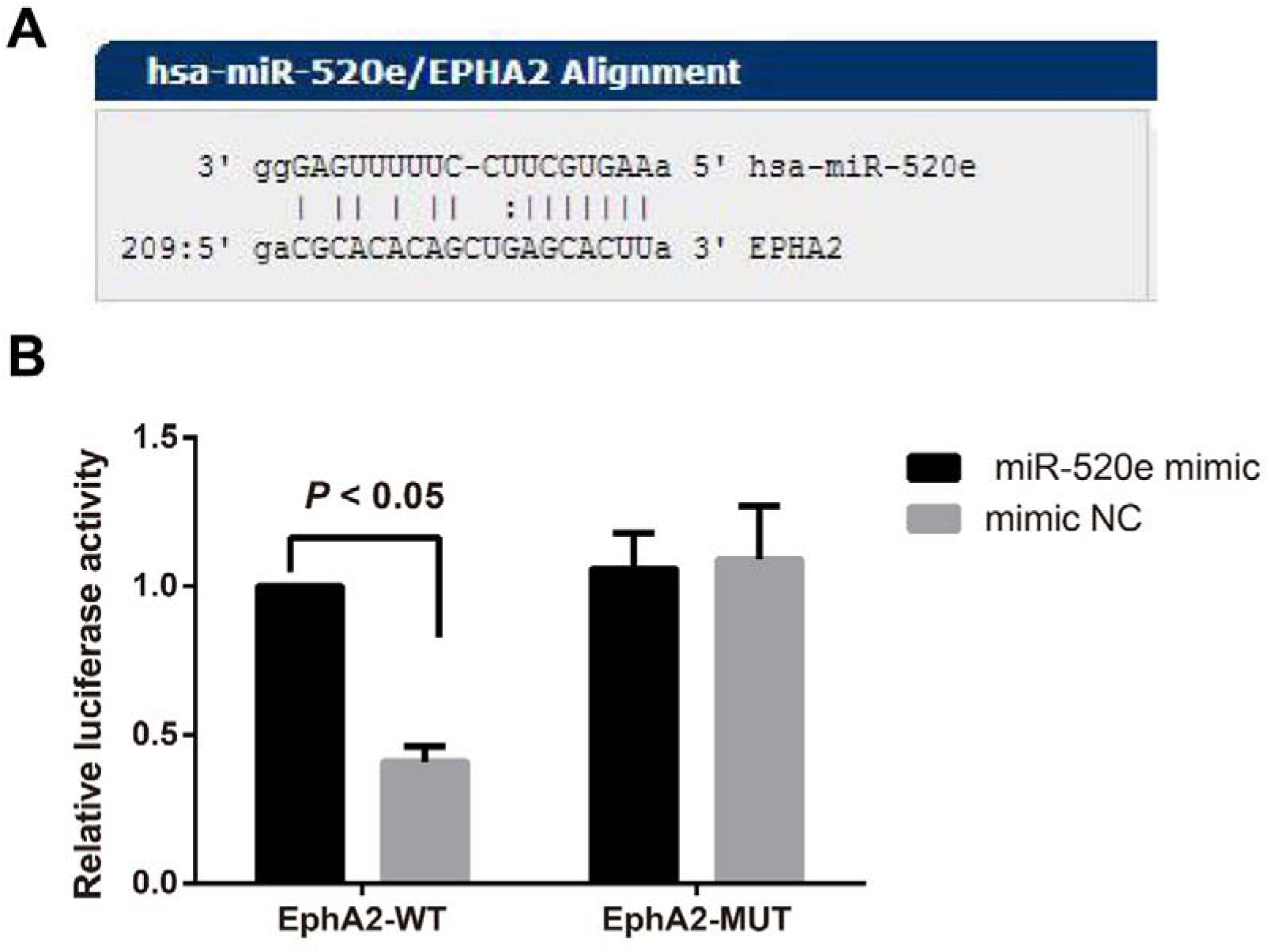
The relative luciferase activity after co-transfection with EphA2 3’UTR and miR-520e mimic. Note: A, Sequence alignments between 3’UTR of EphA2 and miR-520e; B, The relative luciferase activity analysis after co-transfection with EphA2 3’UTR and miR-520e mimic (or miR-520e NC); *, *P* < 0.05 compared with NC group.

### Expression of miR-520e and EphA2 after transfection

After transfection, the expression of miR-520e and EphA2 in HepG2.2.15 and pHBV1.2 + Huh7 cells were tested by qRT-PCR and Western blot. As displayed by the results, NC group and Mock group didn’t statistically differ from each other concerning miR-520e and EphA2 in these two cells (al *P* > 0.05). Compared with Mock group, miR-520e mimic group was increased in miR-520e and decreased in EphA2; however, miR-520e inhibitor and miR-520e inhibitor+ si-EphA2 groups exhibited lower levels of miR-520e, and meanwhile, miR-520e inhibitor group was elevated in EphA2 and si-EphA2 group was down-regulated in EphA2 (all *P* < 0.05). Further, miR-520e inhibitor+ si-EphA2 group showed no observable difference in miR-520e expression (*P* > 0.05), but an obvious reduction in EphA2 protein by comparison with miR-520e inhibitor group (all *P* < 0.05, Figure 4).

**Figure 4.**
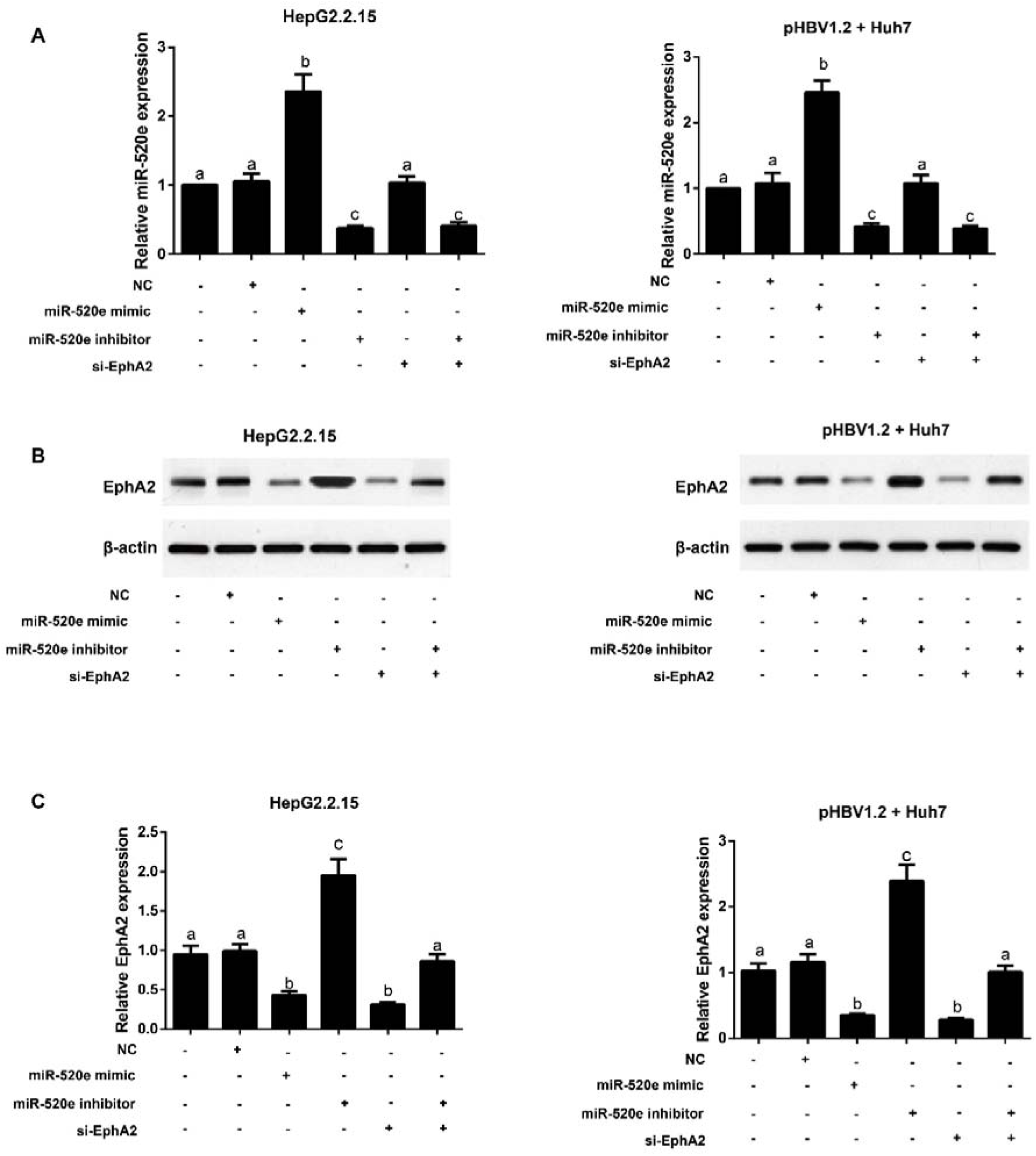
Expression of miR-520e and EphA2 after transfection detected by qRT-PCR and Western blot Note: A, Expression of miR-520e in HepG2.2.15 and pHBV1.2 + Huh7 cells after transfection detected by qRT-PCR; B-C, Protein expression of EphA2 in HepG2.2.15 and pHBV1.2+Huh7 cells after transfection detected by Western blot; The same letters indicated no statistical significance (*P* > 0.05), and the different letters indicated the statistical significance (*P* < 0.05).

### Effect of miR-520e on HBV DNA replication and the levels of HBsAg and HBeAg

As compared with Mock group in both HepG2.2.15 and pHBV1.2 + Huh7 cell lines, miR-520e mimic and si-EphA2 groups were notably reduced, while miR-520e inhibitor group was markedly enhanced in HBV DNA copy number and the levels of HBsAg and HBeAg (all *P* < 0.05, Figure 5), but there was no difference from NC group (all *P* > 0.05). In addition, miR-520e inhibitor +si-EphA2 group was substantially lower than miR-520e inhibitor group, but higher than si-EphA2 group regarding the HBV DNA copy number, HBsAg expression and HBeAg expression (all *P* < 0.05).

**Figure 5.**
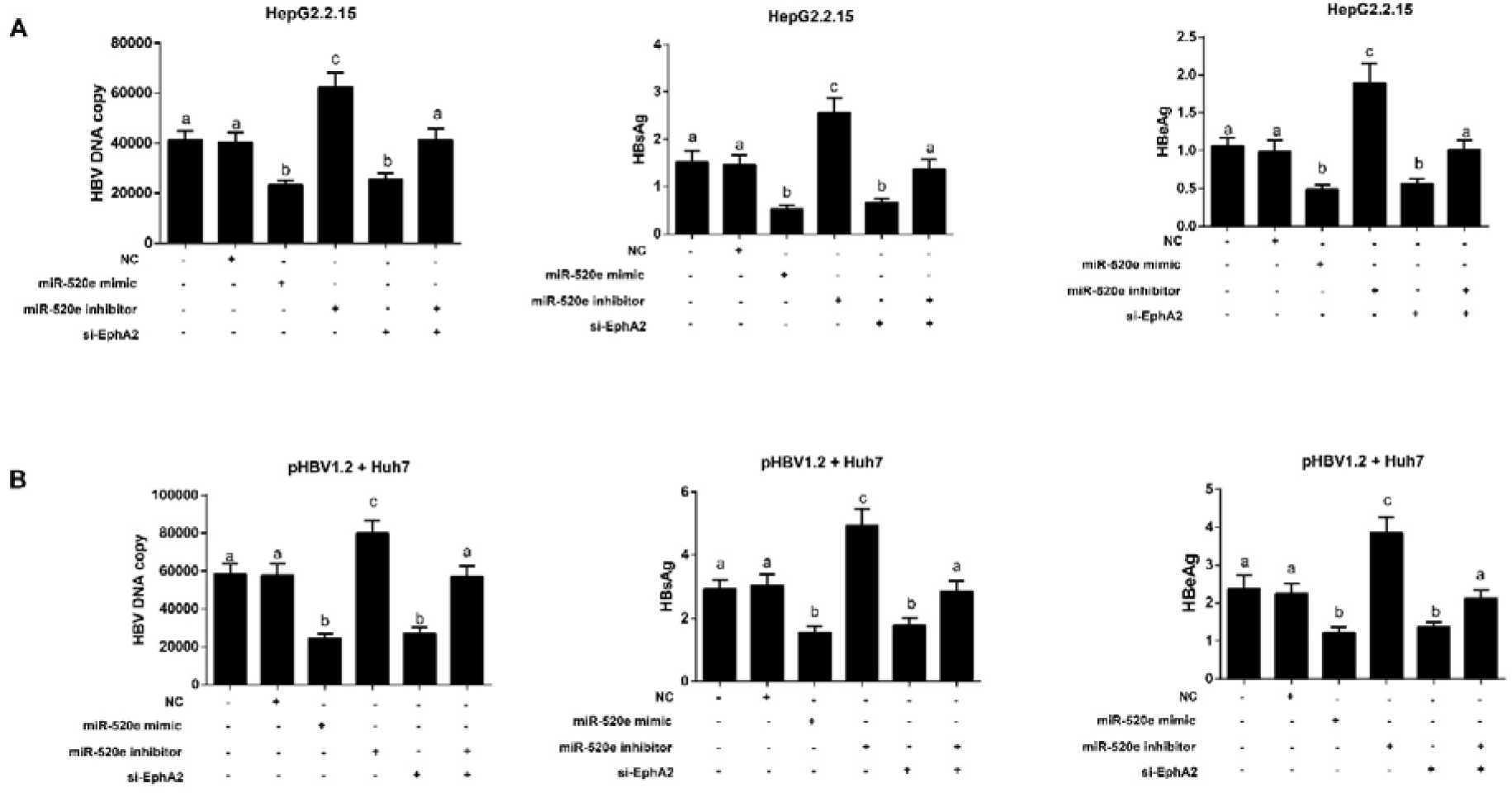
HBV DNA replication and the levels of HBsAg and HBeAg in each group detected by qRT-PCR and ELISA. Note: A, HBV DNA copy number, the levels of HBsAg and HBeAg in each group; B, HBV DNA copy number, the levels of HBsAg and HBeAg in each group; The same letters indicated no statistical significance (*P* > 0.05), and the different letters indicated the statistical significance (*P* < 0.05).

### Cell proliferation after transfection

As illustrated by Figure 6, in HepG2.2.15 and pHBV1.2 + Huh7 cell lines, Mock group and NC group were not statistically different from each other in cell proliferation and apoptosis (*P* > 0.05). Besides, both miR-520e mimic group and si-EphA2 group were significantly reduced in cell proliferation and apparently elevated in cell apoptosis (all *P* < 0.05), but the miR-520e inhibitor group showed the completely opposite trend in these two indexes (all *P* < 0.05). Compared with the miR-520e inhibitor group, the miR-520e inhibitor + si-EphA2 group decreased significantly in cell proliferation and increased remarkably in cell apoptosis (all *P* < 0.05).

**Figure 6.**
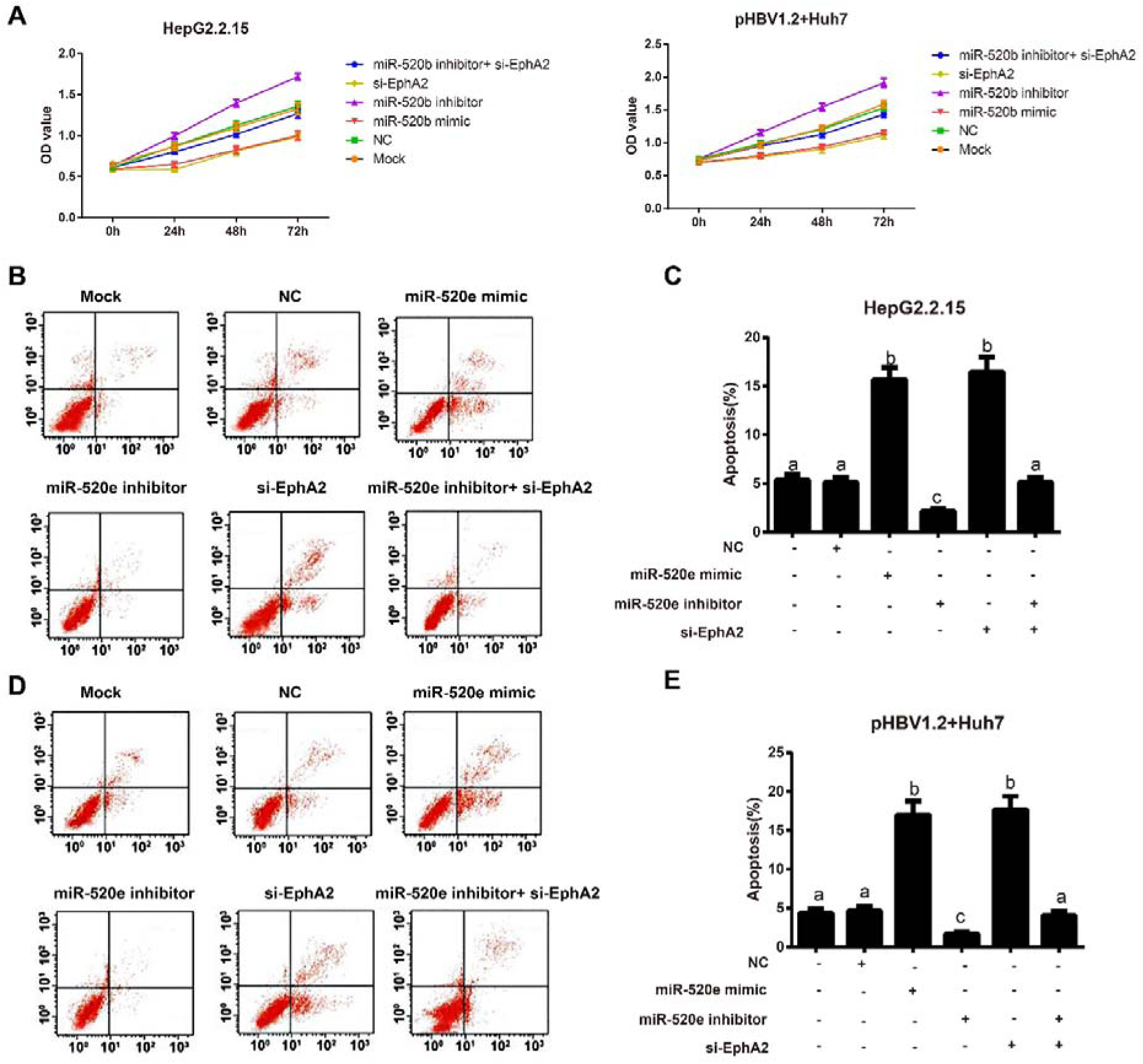
CCK8 and flow cytometry were used to detect the proliferation and apoptosis of HepG2.2.15 and pHBV1.2 + Huh7 cells after transfection. Note: A, The proliferation of HepG2.2.15 and pHBV1.2 + Huh7 cells in each group detected by CCK8 assay; B-C, The apoptosis of HepG2.2.15 cells in each group detected by flow cytometry; D-E: The apoptosis of pHBV1.2 + Huh7 cells in each group detected by flow cytometry; The same letters indicated no statistical significance of inter-group differences (*P* > 0.05), and the different letters indicated the statistical significance (*P* < 0.05).

### Protein expression of p38MAPK and ERK1/2 pathways

The protein expressions of p38MAPK and ERK1/2 pathways in transfected HepG2.2.15 and pHBV1.2 + Huh7 cells were determined by Western blot (Figure 7). Compared with Mock group, miR-520e mimic and si-EphA2 groups were clearly down-regulated, while miR-520e inhibitor group apparently up-regulated in protein expression of p-p38MAPK/p38MAPK and p-ERK1/2/ERK1/2 pathways (all *P* < 0.05). Meanwhile, miR-520e inhibitor group was higher in protein expression of p-p38MAPK/p38MAPK and p-ERK1/2/ERK1/2 pathways than the miR-520e inhibitor + si-EphA2 (all *P* < 0.05).

**Figure 7.**
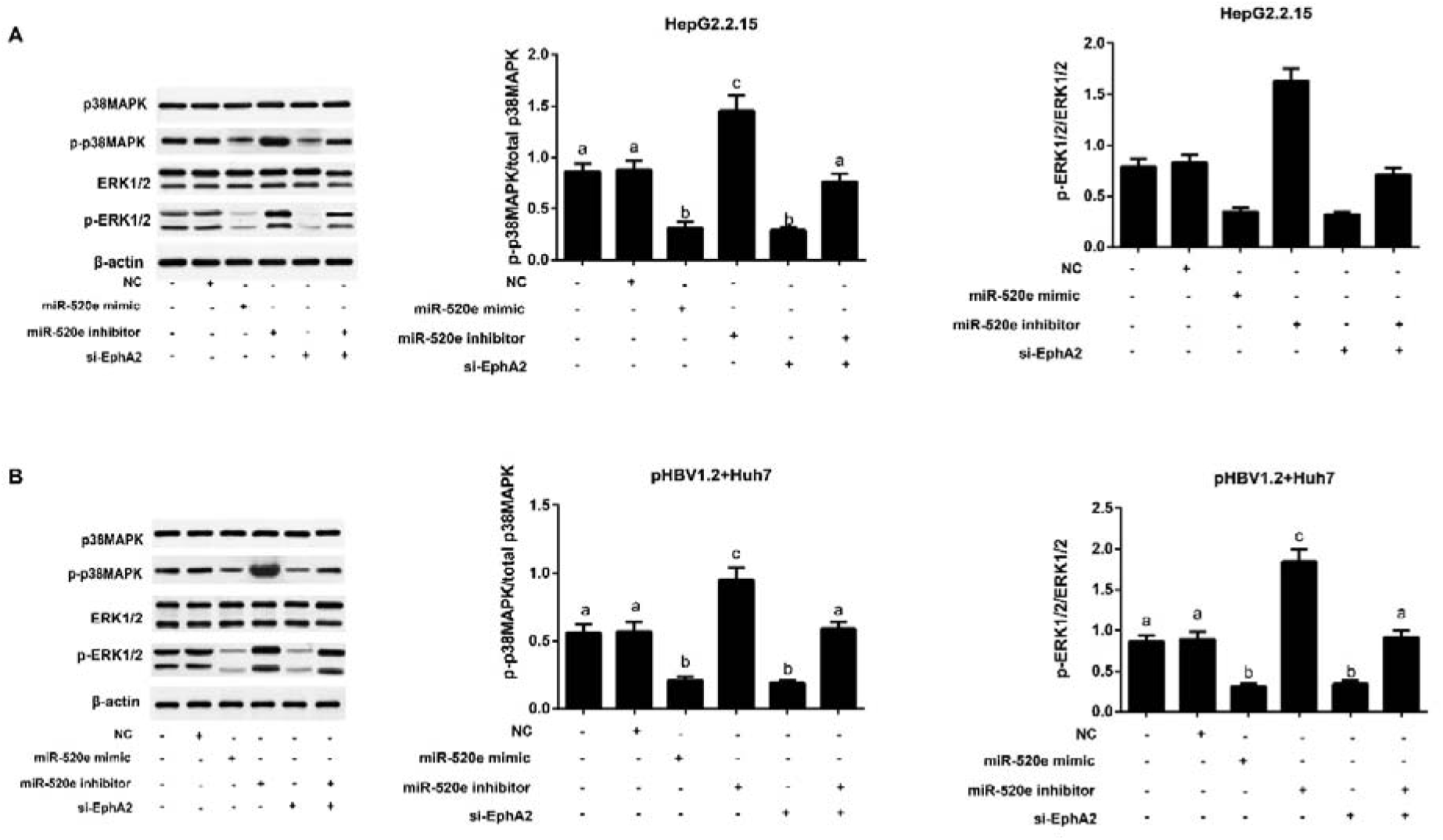
Protein expressions of p38MAPK and ERK1/2 pathways in HepG2.2.15 (A) and pHBV1.2 + Huh7 (B) cells detected by Western blot. Note: The same letters indicated no statistical significance (*P* > 0.05), and the different letters indicated the statistical significance (*P* < 0.05).

### Effect of miR-520e on HBV replication and expression *in vivo*

The levels of miR-520e in liver tissues of HBV-related mice model were detected. According to the results shown in Figure 8, compared with mice in the Normal group, those in the HBV+NC group had statistically decreased expression of miR-520e and apparently increased expression of EphA2 in liver tissues (all *P* < 0.05). At the same time, mice in HBV + miR-520e mimic group were higher in miR-520e and lower in EphA2 than those in HBV + NC group (all *P* < 0.05). On the other hand, mice in HBV + NC group were found to have increased HBV DNA content in serum (all *P* < 0.05), and specifically, mice in HBV+ miR-520e mimic group had lower serum levels of HBV DNA than those in HBV + NC group (all *P* < 0.05).

**Figure 8.**
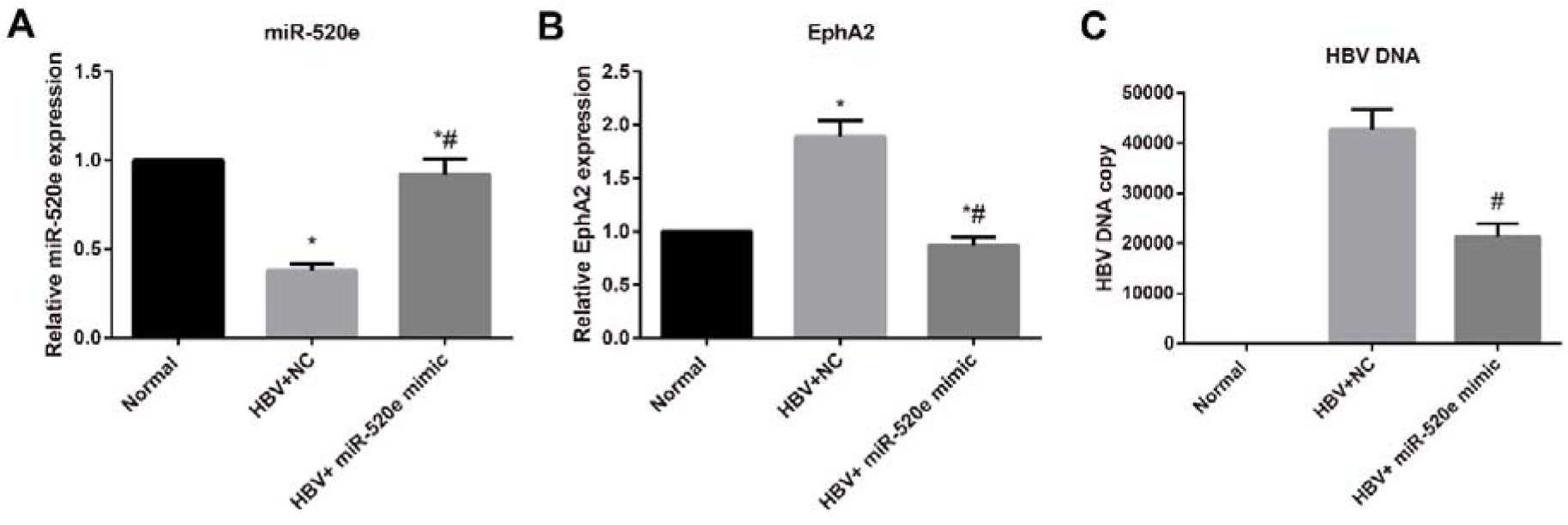
Effect of miR-520e on HBV DNA in rAAV8-1.3HBV-mediated HBV replication model mice Note: A, The levels of miR-520e in liver tissues of mice in each group; B, The expression of EphA2 in liver tissues of mice in each group; C, HBV DNA content in serum of mice in each group; *, *P* < 0.05 compared with Normal group; #, *P* < 0.05 compared with HBV+NC group.

## Discussion

A great number of studies have found the down-regulation of miR-520e, well-documented as a tumor suppressor gene, in a variety of tumor cells, such as breast cancer (Cui et al., 2010), gastric cancer (Li et al., 2016b), lung cancer (Ma et al., 2015). In agreement, this study also discovered a significant reduction of miR-520e in HBV-positive HCC tissues, as well as HCC cell lines with stable HBV replication (like HepG2.2.15 and HepAd38), suggesting that miR-520e was closely related to HBV infection and HCC occurrence. It is worth mentioning that HBx protein, as one of the most important oncogenic proteins for HBV X gene encoding, can regulate the expression of miRNAs at the transcriptional level (Kwon and Cho, 2012), which has been suggested to regulate the HBV virus replication and proliferation to greatly affect the progression of HBV-related HCC (Ng and Lee, 2011; Venard et al., 2000). For example, HBx has previously demonstrated to inhibit miR-15b and miR-205 (Wu et al., 2014; Zhang et al., 2013), or promote miR-224, and thus regulating the growth of HBV-mediated HCC cells (Lan et al., 2014). Therefore, we constructed liver cell line (Huh7-X) and HCC cell line (HepG2-X), both of which HBx was stably expressed, to further investigate the mechanism of miR-520e in HCC. According to the results, the remarkably reduced miR-520e was observed in Huh7-X and HepG2-X cells, but it presented significant increase in a dose-dependent manner in Huh7-X, HepG2-X and HepG2.2.15 cells after interfering HBx, which indirectly confirmed that HBx could down-regulate the expression of miR-520e. Interestingly, Zhang *et al.* also reported that HBx can reduce the expression of miR-520b, a member of miR-520 family, by forming complex with transcription factor Sp-1 and Survivin in liver cells (You et al., 2014; Zhang et al., 2014), which also provided the possibility that the down-regulation of miR-520e in HBV-positive HCC cells may result from the inactivation of miR-520e promoter caused by the interaction between HBx and Sp-1.

In addition, HBV-positive HepG2.2.15 cells and Huh7 cells transfected with pHBV1.2 *in vitro* were selected in this study to explore the role of miR-520e in HBV-mediated HCC. As a result, we found that the over-expressed miR-520e led to the remarkable decrease of HBV DNA copy number and HBV antigen content (including HBsAg and HBeAg), while inhibiting miR-520e contributed to a completely opposite trend, suggesting that miR-520e could effectively inhibit the HBV replication in HCC cells. Similarly, the study of Nicoletta Potenza *et al.* also revealed that miR-125a-5p played the anti-HBV role in HCC cells by interfering the expression of antigen HBsAg (Potenza et al., 2011). Additionally, Zhao *et al.* reported that miR-26b can inhibit viral antigen expression, transcription, and replication, and suppress the activity of HBV enhancer/promoter by down-regulating the target protein (Zhao et al., 2014). In this study, over-expressed miR-520e played its regulatory role in HBV replication by decreasing its target gene EphA2, while silencing EphA2 was found able to reverse the effect of miR-520e inhibitor on promoting HBV replication in HCC cells. To our knowledge, EphA2 belongs to the Eph receptor subfamily and is a member of receptor tyrosine kinases (RTKs) with the characteristics of RTKs (Dunne et al., 2016), which was observed highly expressed in many cancers (Xu et al., 2014), including HCC (Yang et al., 2009). In the study by Hesheng Li *et al.*, the up-regulation of EphA2 was discovered in HCC tissues and cells, and miR-26b could inhibit the proliferation, invasion and migration of HCC cells by targeting EphA2 expression (Li et al., 2015). More importantly, dual-luciferase reporter gene assay in this study identified EphA2 to be the target gene of miR-520e to participate in the progression of HCC. On the other hand, EphA2 has been proved able to affect signal transduction by modulating the expression of many downstream proteins and cytokines (Chang et al., 2008). Further, EphA2 phosphorylation can activate Ras, which combines with GTP to activate Raf and the downstream signaling molecules, such as Src family members, PTKs and PI3K, and the activated Ras-Raf-MAPK signaling pathway can induce cell deformation, adhesion, proliferation, and migration, or inhibit cell apoptosis (Dohn et al., 2001; Macrae et al., 2005; Miura et al., 2009). Notably, HBx has been reported to be the activator of the Src family of tyrosine kinases, which could activate the Ras-Raf-MAPK pathway to affect the proliferation and apoptosis of HBV-related HCC; and at the same time, the activation of Src also enhanced the activity of viral polymerase and thereby promoting HBV replication (Bouchard et al., 2006; Doria et al., 1995; Tarn et al., 2001). As we know, ERK/12 and p38MAPK are two of the most important pathways in MAPK signaling (Bacus et al., 2001), and a previous study confirmed that EphA2 can regulate p38MAPK and ERK1/2 signaling pathway to affect cell growth (Jin et al., 2015). Therefore, we determined the proliferation and apoptosis of HepG2.2.15 and pHBV1.2+Huh7 cell, as well as the expressions of pathway-related proteins, and found that upregulation of miR-520e inhibited cell proliferation and blocked p38MAPK and ERK1/2 pathways, whereas inhibiting miR-520e led to completely opposite results, and si-EphA2 reversed the effect of miR-520e inhibitor. Coincidentally, miR-520d-3p, another member of miR-520 family, could target EphA2 to suppress the proliferation, invasion and migration od gastric cancer cells, as indicated by Li *et al.* (Li et al., 2014), and down-regulation of EphA2 has been previously shown to inhibit Ras/MAPK pathway to inhibit tumor growth (Petty et al., 2012). The above-mentioned findings demonstrated that miR-520e may block the downstream MAPK signaling pathway by targeting EphA2, consequently interrupting the HBV replication and cell proliferation of HBx-mediated HCC. Last but not least, miR-520e can inhibit HBV replication by down-regulating EphA2 was also confirmed by animal models in our experiment.

To sum up, miR-520e was found to be down-regulated in HBV-positive HCC tissues and cells. Besides, over-expression of miR-520e can specifically inhibit EphA2, thus blocking the p38MAPK and ERK1/2 pathways, further resulted in HBV replication and HCC growth. Hence, this study provided a theoretical basis for the prevention and clinical treatment of HBV-related HCC.

## Acknowledgements

We would greatly appreciate the financial support from the National Natural Science Foundation of China (Grants: NSFC, No. 31500148), the Natural Science Foundation of Shandong Province (ZR2012HM082), and the Natural Science Foundation of Shandong Province (ZR2013HQ062).

